# Electromagnetic mapping of the effects of deep brain stimulation and dopaminergic medication on movement-related cortical activity and corticomuscular coherence in Parkinson’s disease

**DOI:** 10.1101/657882

**Authors:** Kousik Sarathy Sridharan, Andreas Højlund, Erik Lisbjerg Johnsen, Niels Aagaard Sunde, Sándor Beniczky, Karen Østergaard

## Abstract

**Background:** Parkinson’s disease (PD) is a debilitating neurodegenerative disorder. PD can be treated with deep brain stimulation (DBS) when dopaminergic medication is no longer a viable option. Both treatments are effective in improving motor symptoms, however, their underlying mechanisms are not fully elucidated yet.

**Objectives:** To study the effects of DBS and dopaminergic medication treatments on cortical processing and corticomuscular drive during movements.

**Methods:** Magnetoencephalography (MEG) was recorded in 10 PD patients and 10 healthy controls, performing phasic hand contractions (hand gripping). Measurements were performed in DBS-treated, untreated and dopaminergic-medicated states; healthy controls received no treatment interventions. Participants performed phasic contractions with their right hand, recorded with electromyography (EMG). Our measures of interest were beta (13-30 Hz) corticomuscular coherence (CMC) and low-gamma (31-45 Hz) power. We used Bayesian statistics on summary values from sensor space data, and we localized the sources of the effects of treatments on beta-CMC and low-gamma power using beamforming.

**Results:** In PD patients, DBS led to reduced CMC values, whereas dopaminergic medication increased beta-CMC values (localized to contralateral M1) to even higher levels than the controls. DBS, on the other hand, increased low-gamma power (localized to M1) compared to controls and to other conditions. Yet both treatments had similar beneficial effects on the patients’ motor symptoms evaluated by UPDRS-III.

**Conclusion:** Despite comparable improvements from both treatments on motor symptoms, DBS and dopaminergic medication seem to have different effects on motor cortical function. This indicates that the treatments undertake different functional strategies to improve PD symptoms.

## Introduction

Parkinson’s disease (PD) is a debilitating neurodegenerative disorder due to the dopaminergic denervation in the striatum and loss of dopaminergic neurons in the substantia nigra pars compacta, in which motor symptoms such as tremor, rigidity, akinesia and gait disturbances manifest as significant disease characteristics (Hughes et al., 1992; Lang and Lozano, 1998; Hammond et al., 2007). Dopamine precursor levodopa remains the primary treatment option. But in advanced PD, motor complications arise due the so-called OFF-states where patients experience motor symptoms at sporadic time periods despite being medicated. In such cases, deep brain stimulation (DBS) of the subthalamic nucleus (STN) is an alternative therapeutic recourse that generally ameliorates the motor complications (Limousin et al., 1998; Østergaard and Aa Sunde, 2006; Johnsen et al., 2009) and is considered more effective in managing symptoms than dopaminergic medication alone (Deuschl et al., 2006).

Both DBS and dopaminergic medication alleviate the motor symptoms, but the underlying mechanisms of DBS, in particular, are still unclear. Several possible mechanisms of DBS have been proposed, and the evidence points to a modulation of the motor cortical network activity as the main driver of symptom improvements (McIntyre et al., 2004; Schnitzler and Gross, 2005; McIntyre and Hahn, 2010). Such a modulation likely happens through antidromic activation of the so-called hyperdirect pathway between M1 and STN (Nambu et al., 2002; Gradinaru et al., 2009; Whitmer et al., 2012).

Outside the basal ganglia network, motor cortex is thus key in understanding the potential differences in effect mechanisms of DBS and dopaminergic medication. In this context, assessing treatment effects on movement related oscillations in the cortical mantle and on corticomuscular communication allow for probing exactly such potential differences.

Corticomuscular coherence (CMC) is known to have influences of both descending and ascending motor pathways (Riddle and Baker, 2005; Baker et al., 2006; Witham et al., 2011) and plays a crucial role in corticomuscular communication (Baker et al., 1997; Salenius et al., 1997; Kilner et al., 2000; van Wijk et al., 2012). Beta-CMC is reduced in untreated PD patients (Salenius et al., 2002). DBS seems to affect beta-CMC variedly with a high degree of inter-subject variability (Park et al., 2009; Airaksinen et al., 2015b) whereas dopaminergic medication has been reported to increment beta-CMC to the level of healthy controls (Salenius et al., 2002).

Motor cortical gamma-band activity (>30 Hz), on the other hand, peaks after movement onset (Muthukumaraswamy, 2010; Litvak et al., 2012) and is sustained throughout the movement (Crone et al., 1998; Muthukumaraswamy, 2010) and is localized to contralateral motor cortex (Crone et al., 1998). The literature on the effects of DBS and dopaminergic medication on motor cortical gamma activity is sparse, however, STN gamma-band activity has shown concomitant increase with improved motor symptoms after levodopa administration (Brown et al., 2001; Kühn et al., 2006).

In order to elucidate the effects of treatments on motor cortical oscillations and CMC, we measured magnetoencephalography (MEG) and electromyography (EMG) from PD patients in DBS-treated, untreated and dopaminergic medicated states while they performed phasic contractions (hand gripping). We used hand gripping instead of an isometric wrist extension task (which is often used in experiments on movement related oscillatory activity) to better exemplify daily activities such as grasping a cup or picking up an object than an isometric task and thereby increase the ecological validity of the target task.

We focused on CMC in the beta band (13-30 Hz), which has been consistently shown for both phasic movements and isometric wrist extensions (Kilner et al., 2000; Marsden et al., 2000; Salenius et al., 2002; Gwin and Ferris, 2012; Pollok et al., 2012; Airaksinen et al., 2015b). Furthermore movement-related low-gamma power (31-45 Hz) was investigated as it has been associated with movement (Crone et al., 1998; Van Wijk et al., 2013) and recently, with increased cortical somatosensory processing during dopaminergic medication (Sridharan et al., 2017). We hypothesized that beta-CMC and low-gamma power in M1 would be modulated by DBS and dopaminergic medication.

## Methods

### Cohort

Twelve PD patients implanted with STN-DBS at Aarhus University Hospital and 10 healthy controls were recruited for the experiment. All patients had been treated with STN-DBS for at least 6 months. Each participant gave their informed consent and the study was approved by the local ethics committee (The Central Denmark Region Committees on Heath Research Ethics). Exclusion criteria consisted of a MMSE score (Mini-Mental State Examination) below 25 and clinically significant depression as assessed by MDI (Major Depression Inventory). A total of two patients were excluded from the study for reasons extraneous to the exclusion criteria: one patient could not tolerate the DBS OFF conditions while the other had a corrupt EMG channel in one condition and hence did not pass the data quality check. The remaining participant cohorts thus consisted of 10 PD patients (age: 60 ±1.7 (SEM); 5 female) and 10 age-matched healthy controls (age: 58.3 ±1.3 (SEM); 3 female). See Tab. 1 for all relevant demographic and clinical details on the two cohorts.

**Tab. 1.**
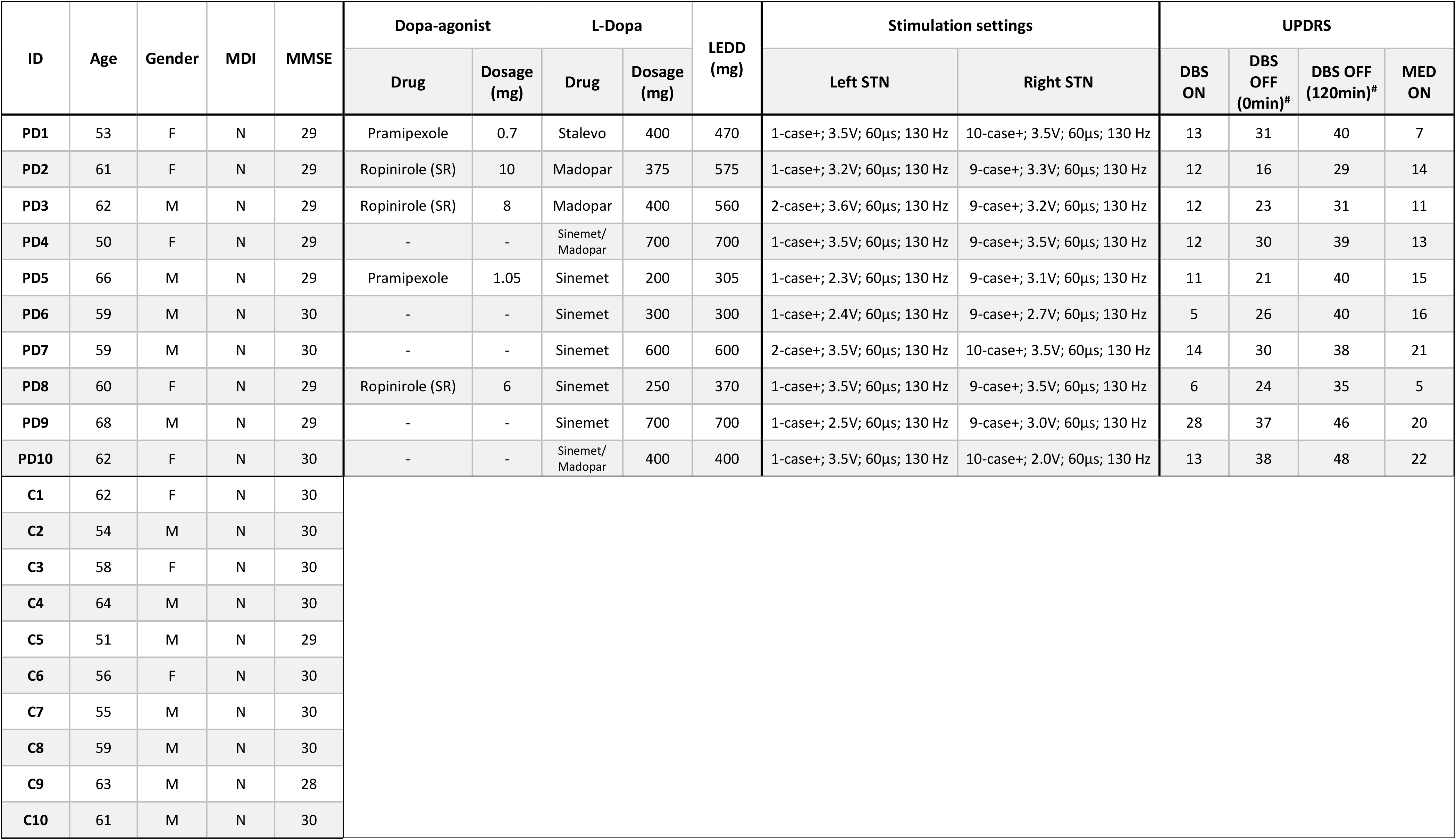
Details of the PD and control cohorts: DBS: deep brain stimulation; MED: L-Dopa medication; UPDRS-III: Unified Parkinson’s Disease Rating Scale (motor part). SR: slow release; LEDD: levodopa equivalent daily dosage; LEDD formula: 100 mg of levodopa = 130 mg of controlled-release levodopa = 70 mg of levodopa + COMT-inhibitor = 1 mg of pramipexole = 5 mg of ropinirole (Mamikonyan et al., 2008). _#_ denotes that the UPDRS-III scoring in those conditions was performed before the experiment session.

### Experimental paradigm

MEG was recorded from participants in supine position with their eyes open. Patients and healthy controls performed self-timed phasic contractions, i.e., grip-relax motions with their right hand, repeatedly over four 30-second sessions with 30-second breaks in-between as shown in Fig. 1. PD patients had ceased taking dopaminergic medication for >12 hours before the experimental sessions. PD patients were recorded in four treatment conditions: DBS-ON, DBS-OFF (0 mins), DBS-OFF (120 mins) and MED-ON conditions. Healthy controls were studied over a similar timescale but without any treatment interventions. Motor symptom assessment (UPDRS-III) was performed before DBS-ON, after DBS-OFF (0 min), after DBS OFF (120 min) to examine the effects of DBS-washout (Temperli et al., 2003) and before the MED ON condition. The intermediary measurements (DBS OFF (30min) through DBS OFF (90 min)) were available only in some patients and hence were not included in the analysis. Participants received breaks between sessions and were taken in and out of the MEG system whenever possible. The experiment was supervised by a movement disorder specialist. One hour prior to the MED ON condition, PD patients were administered 200 mg of levodopa. In order to ensure that the dopaminergic medication had taken effect before performing the final recording session, UPDRS-III assessments were performed prior to the recording session.

**Fig.1.**
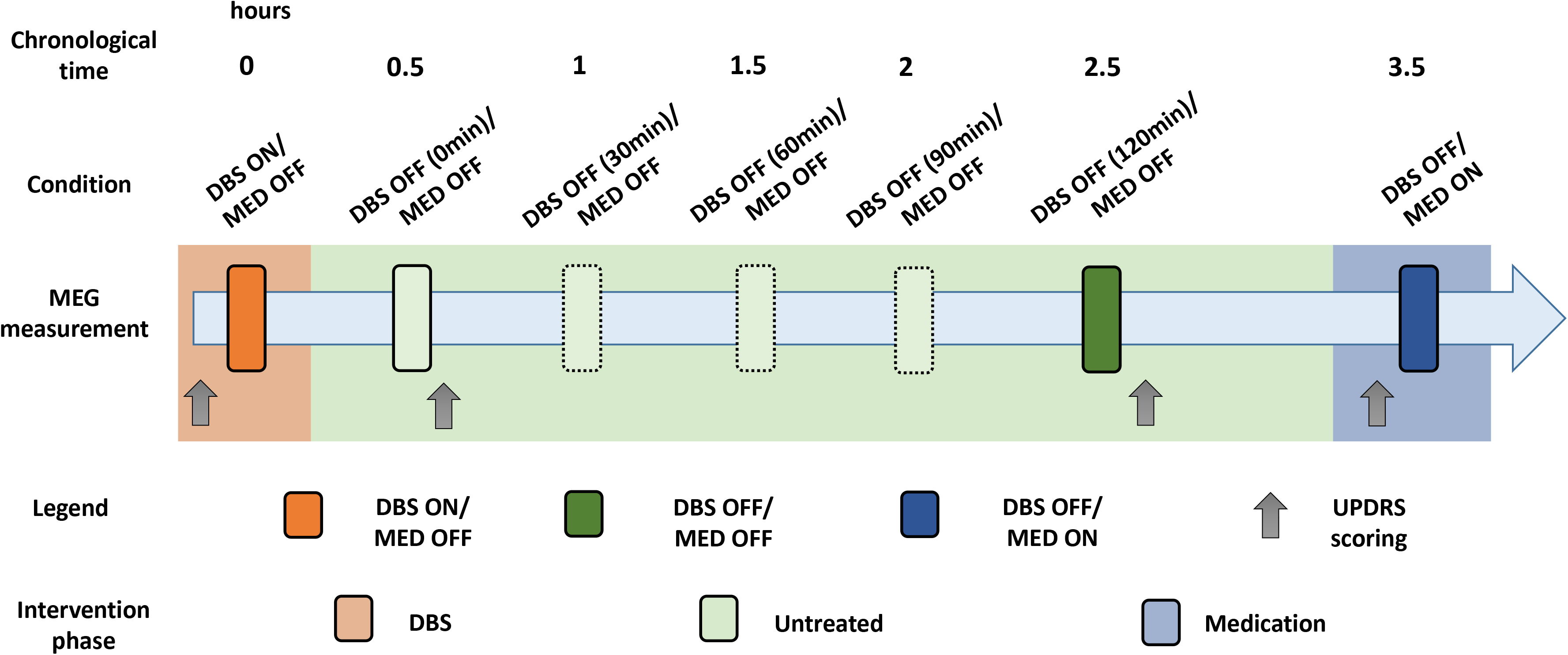
Experimental paradigm: MEG measurements are represented by the vertical bars and each measurement session reflects ~2 minutes of phasic contractions. UPDRS-III was performed at four time points marked by the grey arrows. The background colors represent the three treatment states, namely DBS-treated (orange), untreated (green) and dopaminergic medication-treated (blue) states. Controls did not receive any treatment. Experiment cohorts consisted of 10 PD patients and 10 healthy controls each. See Methods section for more details.

### MEG acquisition

MEG was recorded with a 306-channel Elekta Neuromag TRIUX MEG system placed in a magnetically shielded room. Apart from the neuromagnetic data, vertical electro-oculogram (vEOG), electromyogram (EMG) of the digitorum communis muscle and electrocardiogram (ECG) were also measured. The data were all recorded with a sampling rate of 1 kHz and a passband of 0.03-330 Hz (standard settings for the MEG system).

Head shape was digitized along with the positions of the anatomical fiducials (nasion and right and left pre-auricular points) and the head position indicator coils (HPIs). The anatomical fiducial positions and head shape were then used to co-register the participants’ MEG data and structural MRIs for source localization. The head position measurements in the MEG system were not possible during the DBS ON condition due to interference from the electrical stimulation with the HPIs. We therefore used the head position measurement from the subsequent recording after switching off DBS and before the patient had been taken out of the MEG system (and thus potentially changed his or her position).

### Data preprocessing

The data was first processed using the tSSS algorithm implemented in Maxfilter (Taulu and Simola, 2006) to suppress DBS artefacts and other external magnetic artifacts (subspace correlation limit: 0.9, time window: 10s (Medvedovsky et al., 2009)) and this approach has already been shown to successfully suppress DBS artefacts (Airaksinen et al., 2011, 2012, 2015b, 2015a; Sridharan et al., 2017) and other magnetic interferences (Taulu et al., 2005; Taulu and Simola, 2006; Taulu and Hari, 2009). All data (MEG, EMG, EOG and ECG) were bandpass filtered between 1-100 Hz using a two-pass second order Butterworth filter. An ICA-based artefact rejection routines to suppress ECG- and EOG-related artefacts was implemented using the MNE-Python toolbox (Gramfort et al., 2013). All subsequent analyses of the pre-processed data were carried out using the Fieldtrip toolbox (Oostenveld et al., 2011) as implemented in MATLAB (The MathWorks Inc., Natick, MA, 2000). Data were epoched into 1-second segments around the peaks of the gripping action. To obtain the peaks, EMG data were Hilbert-transformed and the absolute value was computed to obtain the power envelope after which they were standardized. These standardized envelopes were visually inspected to define thresholds for peaks in each condition which were then used to automatically epoch the data. This epoching procedure rendered and average of 91±5 (SEM) epochs in the PD and 97±2 in the control cohort (see Tab. 2 for condition-wise details).

**Tab. 2.** Summary values: Means and SEMs of the differences between measures in each condition and the DBS OFF (120 min) condition, thus used as the reference condition. The SEMs of the group averages in the reference condition are shown in parentheses in that column. The different conditions favored by the statistical analyses are highlighted in bold.

| Measures |  | PD |  |  |  | Control |  |  |  |
| --- | --- | --- | --- | --- | --- | --- | --- | --- | --- |
|  |  | Δ DBS ON | Δ DBS OFF (0min) | DBS OFF (120min) | Δ MED ON | Δ DBS ON | Δ DBS OFF (0min) | DBS OFF (120min) | Δ MED ON |
| Beta-CMC | Mean | -0.012 | 0.001 | **0.094(±0.01)** | **0.030** | -0.024 | -0.020 | **0.110 (±0.025)** | -0.003 |
|  | SEM | 0.008 | 0.007 |  | **0.013** | 0.015 | 0.017 |  | 0.015 |
| Low-gamma Power | Mean | **0.487** | -0.095 | **0.184(±0.15)** | -0.205 | 0.119 | 0.147 | **-0.418 (±0.14)** | 0.079 |
|  | SEM | **0.129** | 0.118 |  | 0.129 | 0.061 | 0.095 |  | 0.106 |
| UPDRS - III | Mean | -26.0 | -11.0 | **38.6 (±1.9)** | -24.2 |  |  |  |  |
|  | SEM | 2.1 | 1.1 |  | 5.5 |  |  |  |  |
| EMG-RMS (mv) | Mean | **0.79** | 0.22 | **0.919 (±0.574)** | **0.58** | -0.02 | 0.08 | **1.73 (±0.259)** | 0.12 |
|  | SEM | **0.20** | 0.35 |  | **0.09** | 0.21 | 0.18 |  | 0.06 |
| Number of trials | Mean | 2 | -11 | **93 (±9)** | 1 | 1 | 1 | **95 (±4)** | 2 |
|  | SEM | 13 | 13 |  | 8 | 4 | 2 |  | 4 |

Reference condition

### Sensor analysis

Power and cross spectral densities (PSD & CSD) between MEG and EMG data were computed using the multitaper method (Mitra and Pesaran, 1999) with discrete prolate spheroidal sequences (DPSS) as tapers, as implemented in Fieldtrip. The central frequencies were 1 to 100 Hz with a bandwidth of 1 Hz. The mean squared coherence was then obtained from the CSD to reflect corticomuscular coherence (CMC). The maxima of the orthogonal gradiometer pairs were used in the case of coherence while the sums of the gradiometer values were used for combining gradiometer pairs for power values. To evaluate the motor output, we computed the mean of the trial-wise root mean square (RMS) which reflects the mean motor output in each condition.

The PSDs were normalized by dividing the PSD spectra by the total power from the range of 1-45 Hz and z-scoring them. We restricted our spectral analysis to below 45 Hz in order not to confound our analyses with potential DBS artefacts from its first sub-harmonics (i.e. at 65-90 Hz depending on individual stimulation frequency). We defined 15 gradiometer pairs in the left (contralateral) sensorimotor area as the sensors of interest (SOI). We defined two a priori frequency bands of interest: beta (13-30 Hz) and low-gamma (31-45 Hz). We inspected any artefactual peaks in the PSD (due to DBS) and dropped those particular frequency bins from further analysis (see Supplementary material, as well as Airaksinen et al., (2012), for a similar approach). We obtained summary values for beta-CMC and low-gamma power by searching for maxima in each of the frequency bands in the SOI. The peak frequencies for each subject, condition and measure were passed on to the source localization procedure.

### Statistical analysis

We performed statistics on patient and control cohorts separately with the control cohort mainly serving to account for potential time-related effects (since counter-balancing the conditions was not possible in our experimental design). We used Bayesian inferencing given by Bayes Factors (BF) to test the effects of treatment on UPDRS-III, EMG-RMS, beta-CMC and low-gamma power. BF is an estimate of relative likelihoods of the observed data under the given hypotheses. We used the *BayesFactor* package (Morey et al., 2015; Sridharan et al., 2017) in R (R Core Team, 2016) to calculate the relevant BFs using the *lmbf* function along with default settings namely, r=0.5 for fixed effects and r=1 (‘nuisance’) for random effects. In each cohort, we estimated the effect of Condition by modelling a null hypothesis (H0) with only subject-specific offsets as random effects against an unconstrained alternative hypothesis (H1) with Condition as fixed effect and subject-specific offsets as random effects. Hypothesis testing moved progressively towards more constraints and comparing the BFs for each of these equality-constrained hypotheses enabled us to obtain meaningful model-driven statistical inferences on the data. We first framed a more constrained hypothesis (H2) to test the three treatment states against each other. We accomplished this by equating DBS OFF (0 min) and DBS OFF (120 min) while allowing DBS ON and MED ON to vary freely. We proceed with testing the specific hypotheses, explained below, only if both H_1_/H_0_ and H_2_/H_0_ showed substantial evidence for the alternative hypothesis.

In order to assess whether only one treatment state stood out from the other two, we tested four specific hypotheses with further equality constraints: (H3) equating conditions DBS ON, DBS OFF (0 min) and DBS OFF (120 min) while allowing MED ON to differ; (H4) equating conditions DBS OFF (0 min), DBS OFF (120 min) and MED ON while allowing DBS ON to differ; (H5) equating DBS ON and MED ON, as well as DBS OFF (0 min) and DBS OFF (120 min) while allowing the two pairs to differ from each other; and (H6) equating DBS ON and MED ON conditions while allowing DBS OFF (0 min) and DBS OFF (120 min) to vary. H1 and H2 in combination with H5 and H6 thus implicitly tested for washout effects of the DBS treatment in that the only difference between the two pairs of hypotheses was that both DBS OFF (0 min) and DBS OFF (120 min) were taken to reflect an undifferentiated untreated state in H2 and H5 in contrast to H1 and H6.

#### Correlations

For both beta-CMC and low-gamma power, we computed their *Kendall’s tau* with both the overall UPDRS-III score and the UPDRS-III bradykinesia (right hand) sub-scores (items: 23 to 25 in the UPDRS scoring manual). We then used the R-function provided along with the recent publication from van Doorn et al. (2016) where the Bayesian inference for correlations in nonparametric data is made on the test statistic in order to obtain BFs. The resulting BFs for *Kendall’s tau* reflect the relative likelihoods of the observed data under the alternative hypothesis (H_1_) that the modelled slope for the relationship between the two variables (e.g. beta-CMC and UPDRS-III) is non-zero against the null hypothesis (H_0_) that the slope is zero. We computed BFs for five correlations: (i, ii) beta-CMC vs UPDRS-III and UPDRS-III bradykinesia (right hand) sub-scores (items: 23 to 25 in the UPDRS scoring manual) in the MED ON condition; (iii, iv) low-gamma power vs. UPDRS-III and UPDRS-III bradykinesia in the DBS ON condition; and (v) EMG-RMS values vs UPDRS-III scores across all conditions.

### Source reconstruction

Based on our observed effects of dopaminergic medication on beta-CMC and of DBS on low-gamma power, we performed source localization only of those two measures in the two treatment conditions, respectively, using beamforming. T1-weighted structural magnetic resonance images (MRI) were acquired for all controls, and we used the pre-operative MRIs for the PD patients. The co-registration of the MEG data and structural MRIs was performed using the fiducials and head shape points recorded during the MEG acquisition.

Each subject’s structural MRI was segmented and the inner skull volume was non-linearly warped to a canonical MRI included in SPM8 (http://www.fil.ion.ucl.ac.uk/spm/software/spm8/) and canonical source model inverse was warped to obtain the subject-specific beamformer grid (spacing of 5 mm) making it comparable across subjects and in correspondence with the specific grid points in the Montreal Neurological Institute (MNI) space (Mattout et al., 2007). The leadfields were computed for each participant and each condition using the single-shell head model based on the individual MRIs of the participants (Nolte, 2003).

Source localization was performed using the Dynamic Imaging of Coherent Sources (DICS) method to localize both beta-CMC and low-gamma power (Gross et al., 2001). In the case of beta-CMC, the recorded EMG channel was used as a reference for the source localization procedure. In the case of low-gamma power, we used the neural activity index (NAI). As already stated, we used the peak frequencies of the beta-CMC and low-gamma power for each subject and condition to focus the source localization. We set the regularization to 5% for beta-CMC and 0% for low-gamma power.

For visualization purposes, we grand-averaged the beta-CMC and low-gamma power source data in the MED ON and DBS ON conditions, respectively, interpolated and plotted them on a canonical volumetric MRI (Colin 27 brain-1mm resolution) with a threshold of 90%. We used the AAL atlas (Automated Anatomical Labelling; Tzourio-Mazoyer et al., 2002), as integrated into the Fieldtrip toolbox, as our anatomical reference and in order to obtain MNI coordinates.

## Results

### UPDRS-III

The UPDRS-III scores of the PD cohort are visualized in Fig 2 (summary values in Tab 2). While evaluating the unconstrained hypothesis (H_1_) we obtained a BF_10_=1.42E12, suggesting that the data were 1.42E12 times more likely to occur under the hypothesis containing Condition as a factor compared to the one without. A BF of this magnitude is considered *very strong evidence* (Kass and Raftery, 1995) and suggests an effect of Condition on the UPDRS-III scores.

**Fig.2.**
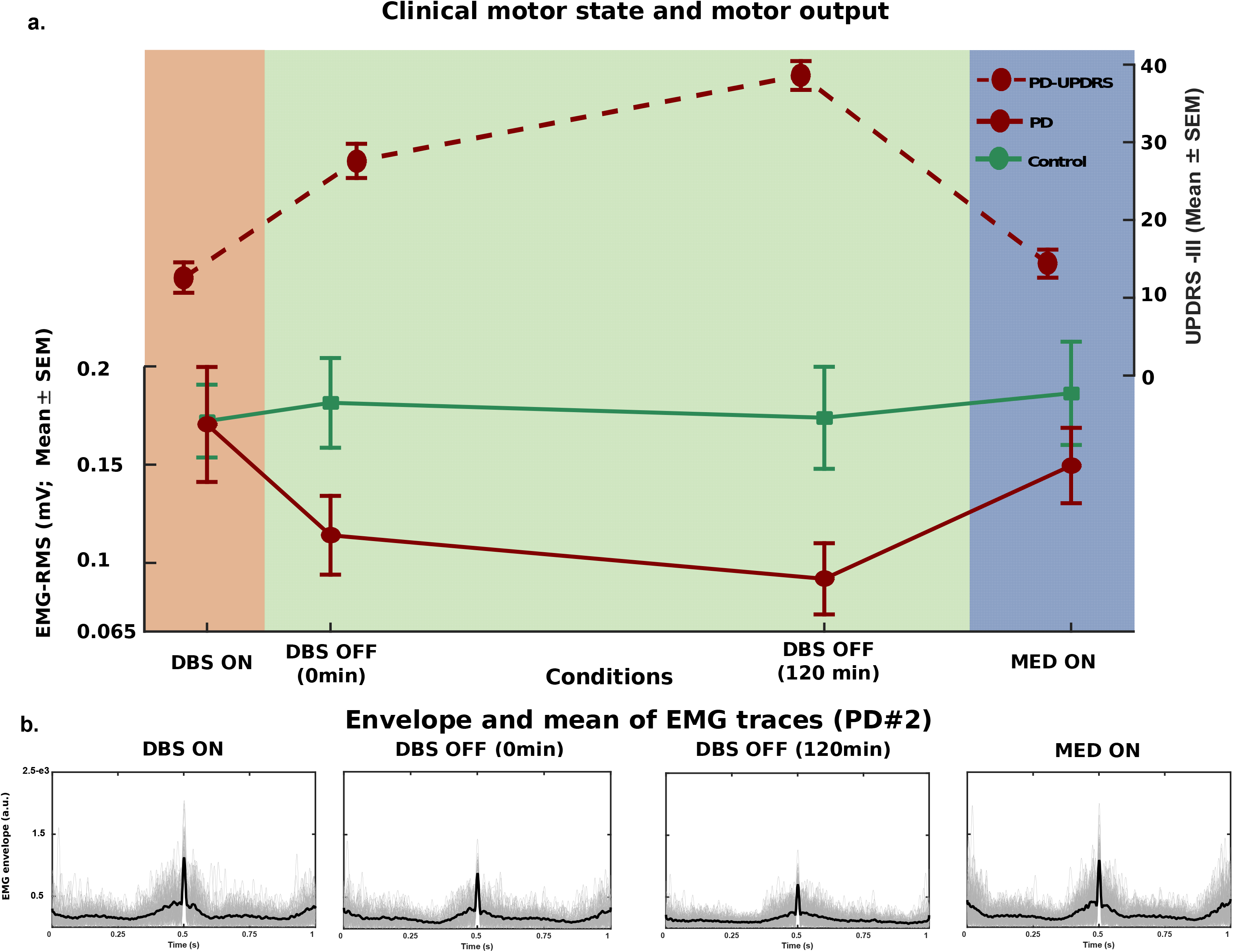
Clinical motor state and motor output: **(a)** Group averages of the UPDRS-III (motor part) scores of the PD cohort are shown at the top (broken maroon line) together with EMG-RMS (motor output) of the PD (solid red) and control cohorts (solid green) at the bottom. Higher values of UPDRS-III scores indicate worsening motor symptoms while higher EMG-RMS values indicate better motor output. While UPDRS-III and EMG-RMS values were similar for both DBS ON and MED ON condition, UPDRS-III scores alone showed a washout effect of DBS (DBS OFF (0 min) compared to DBS OFF (120 min)). See Results section for more details. **(b)** The Hilbert envelope of the epoched EMG trials are plotted along with the trial-averaged envelope (thick line) for each condition from a representative patient (PD#2). See Methods section for more details.

The evaluation of the constrained hypotheses (H_3-6_) revealed highest BF for H_6_ where DBS ON and MED ON conditions were equated while allowing DBS OFF (0 min) and DBS OFF (120 min) conditions to vary. Compared to the unconstrained hypothesis (H_1_), the data were 8.4 times more likely (BF_61_=8.43) to occur under this constrained hypothesis, and 7,375 times more likely (BF_65_=7,375) to occur under this hypothesis compared to the hypothesis (H_5_) with DBS ON and MED ON equated and DBS OFF (0 min) and DBS OFF (120 min) equated. These are considered *positive evidence* and *very strong evidence* (Kass and Raftery, 1995), respectively, and suggest that the two treatments (dopaminergic medication and DBS) lead to similar ameliorating effects on the motor symptoms, while the washout effect of the DBS treatment on the motor symptoms are reflected in the increasing UPDRS-III scores between DBS OFF (0 min) and DBS OFF (120 min) (See Tab.2 for Summary values).

### EMG-RMS

The mean of the EMG-RMS of both cohorts are plotted in Fig. 2 (summary values Tab. 2). Evaluation of the unconstrained hypothesis (H_1_) showed that the EMG-RMS was 177 times (BF_10_=177.38) more likely to arise from this hypothesis with Condition as a factor compared to the hypothesis without (H_0_). This is considered *very strong evidence* (Kass and Raftery, 1995) and suggests an effect of Condition on EMG-RMS.

In evaluation of the specific hypotheses H_3-6_, we obtained the highest BF for H_5_ where DBS ON and MED ON as well as DBS OFF (0 min) and DBS OFF (120 min) were equated. The data were 3.3 times (BF_51_=3.33) more likely to occur under H_5_ compared to the unconstrained hypothesis (H_1_). This is considered *positive evidence* (Kass and Raftery, 1995) and suggests that the PD patients’ EMG-RMS, and hence their motor output, were similar in the two sets of conditions, i.e. in DBS ON and MED ON, as well as in the untreated state with DBS OFF (0 min) and DBS OFF (120 min). The corresponding BFs for Controls showed evidence for the data under the null hypothesis, i.e. for no effects of Condition (see Tab. 3).

**Tab. 3.** Bayes Factors: Bayes Factors for the corresponding alternative hypotheses against the null. Bayes Factors suggesting the strongest evidence for or against the null hypothesis for a given measure are highlighted in bold. ‘1’ denotes DBS ON; ‘2’ denotes DBS OFF (0 min); ‘3’ denotes DBS OFF (120 min); and ‘4’ denotes MED ON. From Kass and Raftery (Kass and Raftery, 1995): BF = 1 to 3 is considered evidence “*barely worth a mention*”; BF = 3 to 20 is considered “*positive evidence*”; BF = 20 to 150 is considered “*strong evidence*”; BF > 150 is considered “*very strong evidence*”.

| BF<sub>x0</sub> -> alternative/null |  | PD |  |  |  |  |  | Ctrl |  |  |  |  |  |
| --- | --- | --- | --- | --- | --- | --- | --- | --- | --- | --- | --- | --- | --- |
|  |  | All conditions<br>(H<sub>1</sub>/H<sub>0</sub>) | 1 ≠ 2=3 ≠ 4<br>(H<sub>2</sub>/H<sub>0</sub>) | Specific hypotheses |  |  |  | All conditions<br>(H<sub>1</sub>/H<sub>0</sub>) | 1 ≠ 2=3 ≠ 4<br>(H<sub>2</sub>/H<sub>0</sub>) | Specific hypotheses |  |  |  |
|  |  |  |  | 1:3 ≠ 4<br>(H<sub>3</sub>/H<sub>0</sub>) | 1 ≠ 2:4<br>(H<sub>4</sub>/H<sub>0</sub>) | 1=4 ≠ 2=3<br>(H<sub>5</sub>/H<sub>0</sub>) | 1=4 ≠ 2≠3<br>(H<sub>6</sub>/H<sub>0</sub>) |  |  | 1:3 ≠ 4<br>(H<sub>3</sub>/H<sub>0</sub>) | 1 ≠ 2:4<br>(H<sub>4</sub>/H<sub>0</sub>) | 1=4 ≠ 2=3<br>(H<sub>5</sub>/H<sub>0</sub>) | 1=4 ≠ 2≠3<br>(H<sub>6</sub>/H<sub>0</sub>) |
| Beta-Left hemisphere | CMC | 3.4503 | 6.8717 | 14.2298 | 1.4248 | 0.4017 | 0.2402 | 0.4817 |  |  |  |  |  |
|  | Power | 0.7007 |  |  |  |  |  | 0.1455 |  |  |  |  |  |
| Low-gamma Left hemisphere | CMC | 0.5995 |  |  |  |  |  | 2.5996 |  |  |  |  |  |
|  | Power | 1253.1 | 3104.683 | 3.3677 | 6727.95 | 0.7903 | 0.4677 | 0.4119 |  |  |  |  |  |
| EMG-RMS |  | 177.38 | 318.90 | 0.6835 | 16.82 | 590.08 | 350.98 | 0.1631 |  |  |  |  |  |
| UPDRS-III |  | 1.40E+12 | 2.80E+08 | 9.5999 | 65.222 | 1.60E+09 | 1.18E+13 |  |  |  |  |  |  |

### Sensor analysis

#### CMC

The group means and the grand averaged topographies of beta-CMC are shown in Fig. 3a. The grand averages of the beta-CMC showed relatively focal effects in a few sensors over the contralateral sensorimotor area (See Fig. 3b). The grand-averaged coherence spectra based on the sensor that captured the maximal beta-CMC is shown in Fig. 3c. The grand-averaged spectra and the spectra at the individual level showed no artefactual peaks due to DBS. Evaluating the effects of treatments on beta-CMC revealed that the beta-CMC data were 3.5 times (BF_10_=3.45) more likely to arise from the hypothesis containing Condition as a factor and specified as the three treatment states (H_1_) compared to the null hypothesis without Condition as a factor (H_0_). This is considered *positive evidence* (Kass and Raftery, 1995) suggesting an effect of treatment on beta-CMC.

**Fig.3.**
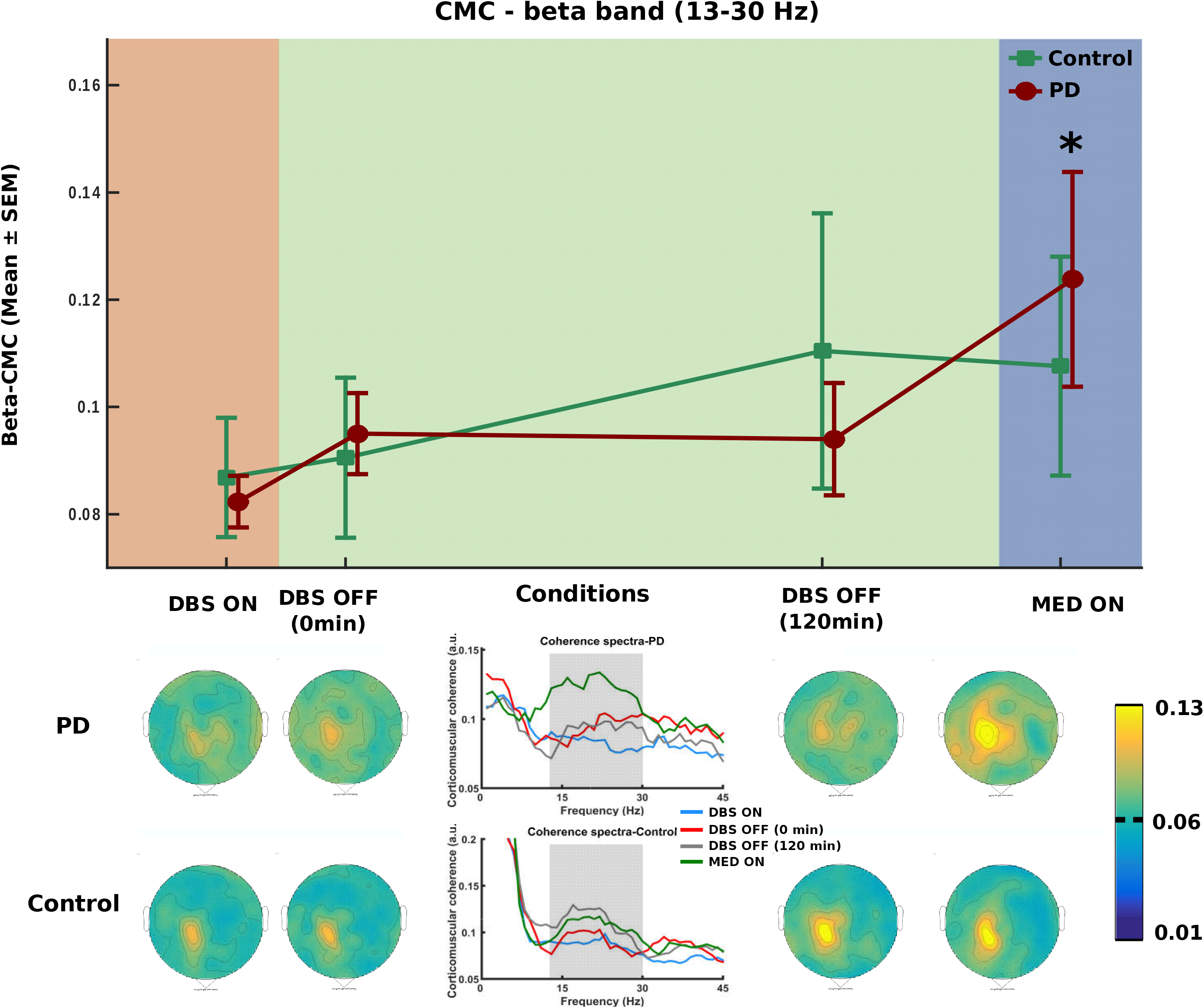
Beta-CMC – trends and topographies: **(a)** Changes in beta-CMC during phasic contractions in the PD (red line) and control (green line) cohorts. The * sign denotes the condition that was found to differ substantially from the others in the PD cohort according to our statistical analyses. Errorbars reflect ±SEM. **(b)** Topographic plots of the group-averaged condition-wise beta-CMC for the PD (top row) and control (bottom row) cohorts. Source localization confirmed that the beta-CMC during MED ON was localized to the contralateral sensorimotor area (see Fig. 5). See Methods section for more details. **(c)** The grand averaged coherence spectra based on the sensor which captured the maximal coherence in each subject and condition are shown.

On evaluating the specific hypotheses (H_3-6_), we found the highest BF for H_3_ where DBS ON, DBS OFF (0 min), DBS OFF (120 min) are equated and only MED ON is allowed to differ. The beta-CMC data were 14.2 times (BF_30_=14.23) more likely to arise from this hypothesis compared to the null hypothesis (H_0_) and 2 times (BF_32_=2.06) more likely to arise from the specific hypothesis H_3_ compared to the constrained hypothesis (H_2_). Both BF_30_ and BF_20_ are considered *positive evidence* (Kass and Raftery, 1995) and since BF_30_ is the higher of the two, it indicates that the MED ON condition was different from the other three conditions and thus that dopaminergic medication affects beta-CMC. The corresponding BFs for the Controls showed evidence for the null hypothesis, i.e., no effects of Condition (or Time, see Tab. 3).

#### Power

Fig. 4a shows the group means and grand-averaged topographies of low-gamma power. In the DBS ON condition in the PD patients, the low-gamma power showed a focus in activity in sensors over the contralateral sensorimotor area (Fig. 4b), quite similar in distribution to the beta-CMC. The grand-averaged power spectra based on the sensor that captured the maximal low-gamma power is shown in Fig. 4c. Individual power spectra showed subject-specific artefactual frequency bins within the frequency range of interest, which were dropped from the analysis (see Methods). Evaluating the effect of treatment on low-gamma power estimated that the data were 3,105 times (BF_20_=3,104.68) more likely to occur under the hypothesis with Condition as a factor and with the DBS OFF (0 min) and DBS OFF (120 min) equated (H_2_) compared to the null hypothesis. This is considered *very strong evidence* (Kass and Raftery, 1995) suggesting an effect of treatment on low-gamma power. Evaluation of the specific hypotheses (H_3-6_) revealed that the low-gamma power data were 6728 times (BF_40_=6727.95) more likely to occur under the hypothesis where DBS OFF (0 min), DBS OFF (120 min) and MED ON are equated (H_4_) compared to the null hypothesis (H_0_). This is considered *very strong evidence* (Kass and Raftery, 1995) and suggests that the low-gamma power is different in the DBS-treated condition compared to the other three conditions and thus an effect of DBS on movement-related low-gamma power. The corresponding BFs for the Controls showed evidence for the null hypothesis and thus suggest no effects of Condition (or Time, see Tab. 3).

**Fig.4.**
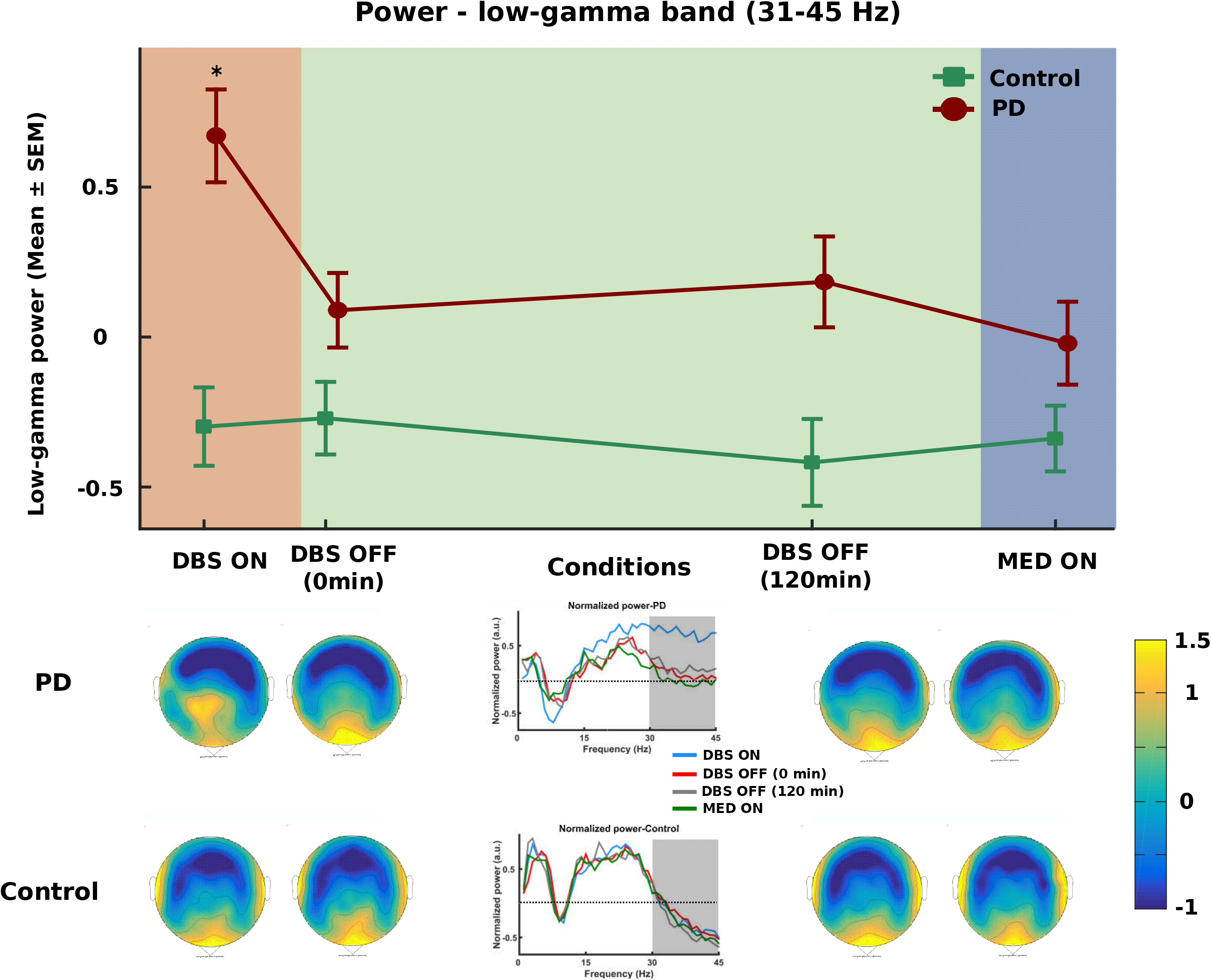
Low-gamma power – trends and topographies: **(a)** Changes in low-gamma power during phasic contractions for the PD (red line) and control (green line) cohorts. The * sign denotes the condition that was found to be different substantially from the others in the PD cohort according to our statistical analyses. Errorbars reflect ±SEM. **(b)** Topographic plots of the group-averaged condition-wise low-gamma power for the PD (top row) and control (bottom row) cohorts. Source localization confirmed that the low-gamma power during DBS ON was localized to the contralateral sensorimotor area (see Fig. 5). **(c)** The grand averaged power spectra based on the sensor which captured the maximal low-gamma power in each subject and condition are shown.

In order to rule out any potential effects in the DBS ON condition due to confounds from sub-harmonics and interference from the electrical stimulation (at 130 Hz), we conducted a parallel sham-analysis for low-gamma power with 15 gradiometer pairs over the right (ipsilateral) hemisphere and with a set of 8 gradiometer pairs over the occipital region. Low-gamma summary values obtained from the right and occipital SOIs did not show any modulation by the treatments, thereby confirming that the increase in low-gamma power in DBS ON over the sensorimotor area is very unlikely to be an artefact of the electrical stimulation (see Supplementary material). We also correlated the DBS voltage levels for the patients’ left and right hemispheres with their observed low-gamma power over the contralateral sensorimotor area as a further test for potential confounds from the electrical stimulation (See Correlations).

### Correlations

UPDRS-bradykinesia (right-hand) sub-scores correlated with the beta-CMC in the MED ON condition (BF_10_=3.83; τ=-0.57), suggesting that the observed data were 3.8 times more likely to occur under the alternative (H_1_) than under the null hypothesis (H_0_). The overall UPDRS-III scores did not show any substantial evidence for the data under the alternative or the null hypothesis (BF_10_=1.86; τ=-0.47). While the former BF (UPDRS-bradykinesia) is considered *positive evidence*, the latter BF (UPDRS-III) denotes *evidence barely worth a mention* (Kass and Raftery, 1995). The low-gamma power was not correlated with the overall UPDRS-III scores (BF_10_=0.63; τ=-0.26) or the UPDRS-bradykinesia (right-hand) sub-scores (BF_10_=0.59; τ=0.24). The BFs suggest that the observed relationships between the low-gamma power and the UPDRS-III and UPDRS-bradykinesia scores were 1.6 and 1.7 times more likely to occur under the null hypothesis (H_0_) with zero slope, respectively. Neither BFs are considered more than *evidence barely worth a mention* (Kass and Raftery, 1995).

The UPDRS-III scores correlated with the motor output as measured by EMG-RMS (BF_10_=132.75; τ =-0.41). The BF suggests that the observed data were 133 times more likely to occur under the alternative hypothesis (H_1_) with non-zero slope than under the null hypothesis (H_0_) with zero slope. This is considered *very strong evidence* (Kass and Raftery, 1995).

For the potentially confounding correlations between DBS voltage levels and low-gamma power, we did not find any substantial evidence for or against the data under the null or the alternative hypotheses. When modelling the correlations between DBS voltage and low-gamma power in the DBS ON condition, the BFs suggested weak evidence for the data under the null hypothesis (H_0_) (left: BF_10_=0.39, τ=0.03; right: BF_10_=0.48, τ=-0.17). The low-gamma power results thus do not seem to be confounded by the electrical stimulation, albeit the correlation analyses only provide weak evidence for this conclusion (see Supplementary material).

### Source reconstruction

Fig. 5 shows the grand-averaged source localization maps of the beta-CMC and low-gamma power in the PD cohort in the MED ON and DBS ON conditions, respectively. The maxima of the cortical source were localized to the precentral gyrus (motor cortex) in the left hemisphere, contralateral to the active muscular movement for both metrics (MNI coordinates of the peaks of the grand averaged source estimates: beta-CMC in MED ON condition: x=-32; y=0; z=44; low-gamma power in DBS ON condition: x=-32; y=-18; z=58).

**Fig.5.**
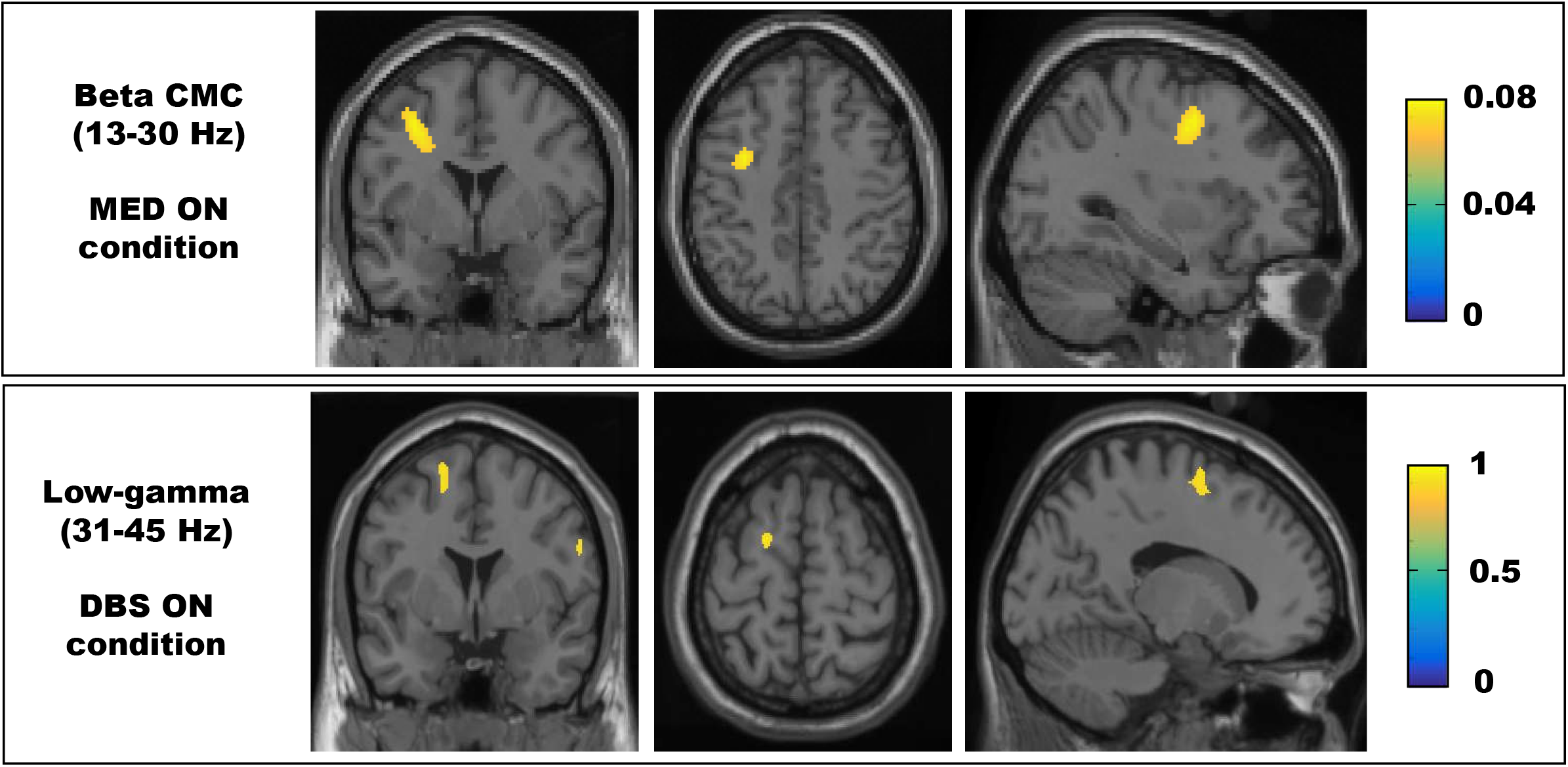
Source estimates of beta-CMC and low-gamma power: Group-averaged source estimates of the beta-CMC (top row) in the MED ON condition and of the low-gamma power (bottom row) in the DBS ON condition. Low-gamma power was localized to contralateral M1.

## Discussion

We have shown in this study, differentiated effects of DBS and dopaminergic medication on movement related oscillations in the motor cortex and the corticomuscular interaction while both treatments showing similar ameliorating effects on the motor symptoms in PD patients. Dopaminergic medication augments beta-CMC (13-30 Hz) in PD patients (localized to contralateral precentral gyrus) while DBS increases low-gamma (31-45 Hz) spectral power (also localized to contralateral precentral gyrus).

### Mechanism of dopaminergic medication on beta-CMC and motor symptoms

The beta-CMC in the medicated state was correlated with improvements in the UPDRS-III scores and the UPDRS-bradykinesia (right hand) sub-scores, indicating a possible link between the general improvement of motor symptoms through dopaminergic medication and an increase in beta-CMC. Studies with healthy subjects have shown that better performance with fewer errors are accompanied by increased beta-CMC (Kristeva et al., 2007; Witte et al., 2007; van Wijk et al., 2012) indicating that higher CMC may lead to better compliance and performance.

While the literature on DBS effects on beta-CMC have shown mixed effects with some indicating a significant increase in beta-CMC due to DBS (Park et al., 2009) and others reporting no group effect of DBS on beta-CMC, pointing to high inter-individual differences in the effect of DBS (Airaksinen et al., 2015b). Dopaminergic medication seems to increase beta-CMC to the levels of healthy controls (Salenius et al., 2002), though Hirschmann et al. (2013) report no significant effects of dopaminergic medication on beta-CMC. Our results are thus in line with the increased beta-CMC during MED ON reported by Salenius et al. (Salenius et al., 2002), though in comparison with healthy controls we report an overshoot in the beta-CMC. Such differences between the two studies may arise from differences in the nature of the tasks applied. While Salenius et al. (Salenius et al., 2002) used an isometric contraction task, we used a phasic hand gripping task, as it has been shown that CMC varies depending on the nature of the task (Kilner et al., 2000). Our results are contrary to results which show beta-CMC is not affected by dopaminergic medication (Hirschmann et al., 2013), it is important to note that the experiment in said report were performed with subcutaneous apomorphine medication, and although it has a very short half-life and was stopped two hours before experiment, residual subcutaneous apomorphine may have confounded the finding. So we compliment the results previously reported by eradicating the potentially confounding effect of concomitant medication and extend the results already reported by Salenius et al. (2002) by incorporating a DBS ON condition.

In rat models, dopamine has been shown to affect local circuits and the cortical interneuron activity which in turn affects the pyramidal neuron activity (Gao and Goldman-Rakic, 2003; Gao et al., 2003). Dopaminergic medication could influence the direct and indirect pathways in the cortico-basal ganglia circuitry by binding to the dopaminergic receptors (D1 and D2) in striatum (Surmeier et al., 2007). These two results are indicative of the influence of dopamine on striatal input to the basal ganglia and pyramidal-interneuron interplay and eventually on the corticomuscular drive. Moreover, beta-CMC increases have been shown to improve performance (Kristeva et al., 2007; Witte et al., 2007), and the increase we observe may thus be a compensatory mechanism to improve the recalibration process and reduce errors during movements as part of the overall improvement in motor action caused by dopaminergic medication in PD patients.

### Mechanism of DBS on low-gamma power and motor symptoms

DBS, unlike dopaminergic medication, seems to amplify movement-related low-gamma (31-45 Hz) power over the contralateral motor cortex and DBS also improved motor symptoms. Low-gamma oscillations have been shown to be somatotopically specific to areas in the M1 and to coincide with maps generated with cortical electrical stimulation (Crone et al., 1998). It has also been shown that gamma activity facilitates movement (Brown et al., 2001; Kühn et al., 2006; Cheyne and Ferrari, 2013) and that dopaminergic medication increases gamma activity in the STN in combination with improvement of motor symptoms suggesting a potential role for gamma activity in the basal ganglia-cortical loop (Brown et al., 2001; Kühn et al., 2006). Increased activity in the low-gamma band is indicative of the activity of the underlying neuronal population (Manning et al., 2009; Miller et al., 2009) and the increased low-gamma band activity during DBS ON might indicate a higher level of activation of the motor cortex during movement (Muthukumaraswamy, 2010).

We speculate that our reported effect of STN-DBS on movement-related low-gamma power arise through antidromic activation of a “hyperdirect” pathway between M1 and STN (Nambu et al., 2002; Gradinaru et al., 2009; Nambu and Tachibana, 2014). Substantial projections from deep cortical layers to the ipsilateral dorsolateral area of STN have been found in monkey (Nambu et al., 2000) and in humans (Fernandez-Miranda et al., 2012; Petersen et al., 2017), supporting the existence of such a hyperdirect pathway. The central role of the antidromic activation of the hyperdirect pathway between M1 and STN in the therapeutic effect of DBS has been made plausible through optogenetic studies (Gradinaru et al., 2009) and computational studies (Kang and Lowery, 2014). It has also been shown that cortical oscillatory activity is functionally coupled to STN oscillations through phase-amplitude coupling, which is in turn affected by DBS (Whitmer et al., 2012; de Hemptinne et al., 2013; Shimamoto et al., 2013). Hence, DBS potentially modulates the cortical low-gamma oscillations through its effects on the basal ganglia-cortical functional network via the “hyperdirect pathway”.

However, we note that the present study was performed with a modest cohort of 10 PD patients and 10 healthy controls, and the evidence for our observed treatment effects on movement related cortical oscillations are moderate in nature and thus necessitate studies with larger cohorts to ensure the potential reproducibility of the results.

In conclusion, we have shown differentiated effects of DBS and dopaminergic medication on cortical activity during phasic hand contractions. While both treatments affect motor symptoms similarly, each of the treatments may employ different functional modulation strategies through differentiated anatomical pathways in their improvement of the motor output.

## Supporting information

Supp. Fig. S1: DBS artefact rejection

Supp. Fig. S2: Correlation between beta-CMC and clinical motor state

Supp. Fig. S3: Influence of DBS artefacts on low-gamma power

Supp. Tab. ST1: Bayes Factors for the sham analyses

## Acknowledgements

We thank our PD patients and healthy controls who participated in the study. We would like to thank Sarang Dalal for his advice on the source analysis, Lau Møller Andersen and Mads Jensen for their advice on the MEG pre-processing, and Jonas Kristoffer Lindeløv for his advice on the statistical analyses. This study is based on work that has been funded by the Danish Research Council for Independent Research, the Danish Parkinson Association, Central Denmark Research Foundation, Aarhus University and Aarhus University Hospital.

## Supplementary material

**Supp. Fig. S1: DBS artefact rejection:** Power spectra of a sample PD patient (PD#4) during DBS ON and MED ON conditions from three SOIs (left and right hemispheres and the occipital area). Bins showing sharp peaks occurring in all three SOIs (here at 34 Hz) were considered artefactual and dropped from further analysis (see Methods).

**Supp. Fig. S2: Correlation between beta-CMC and clinical motor state:** Scatter plots of the beta-CMC and UPDRS-III (left panel) and UPDRS-III-hypokinesia sub-scores from the right hand (right panel) in the MED ON condition. The corresponding BFs and the estimated correlation coefficients are in the titles of the respective plots. While there is *positive evidence* for a negative correlation between beta-CMC and the UPDRS-III-bradykinesia sub-scores, the evidence in the case of UPDRS-III scores is *barely worth a mention*. See Correlations section for more details.

**Supp. Fig. S3: Influence of DBS artefacts on low-gamma power: (a)** Scatter plots of the low-gamma power and the DBS voltages in the left (left panel) and right (right panel) hemispheres. The corresponding BFs and the estimated correlation coefficients are in the titles of the respective plots. While both BFs denote *evidence barely worth a mention* for zero-slope, indicating no substantial evidence for the data under the null hypothesis, there is even less substantial evidence for the data under the alternative hypothesis. **(b)** Low-gamma power in the occipital region across the four conditions. The plot shows no substantial differences in low-gamma power in the occipital region between treatments. This suggests that the increased low-gamma power in the DBS ON condition is a localized effect over the sensorimotor area and that it is not due to corruption from sub-harmonics of the DBS artefacts.

**Supp. Tab. ST1: Bayes Factors for the sham analyses:** Bayes Factors for the corresponding alternative hypotheses against the null for the sham SOIs, namely, the right hemisphere and the occipital area. Evaluation of the hypothesis (H_1_) revealed that the low-gamma power data was more likely to occur under the null hypothesis compared to the alternative hypothesis with Conditions as factor (H_1_). This is considered *positive evidence* (Kass and Raftery, 1995) and suggests that the low-gamma power was not substantially different between conditions and thus no noticeable effect of DBS treatment on movement-related low-gamma power over the right hemisphere and over the occipital area. This confirms that the increase in low-gamma power in DBS ON over the left sensorimotor area is very unlikely to be an artefact of the electrical stimulation.

## References

Airaksinen K, Butorina A, Pekkonen E, Nurminen J, Taulu S, Ahonen A, Schnitzler A, Mäkelä JP (2012) Somatomotor mu rhythm amplitude correlates with rigidity during deep brain stimulation in Parkinsonian patients. Clin Neurophysiol 123:2010–2017.

Airaksinen K, Lehti T, Nurminen J, Luoma J, Helle L, Taulu S, Pekkonen E, Mäkelä JP (2015a) Cortico-muscular coherence parallels coherence of postural tremor and MEG during static muscle contraction. Neurosci Lett 602:22–26.

Airaksinen K, Mäkelä JP, Nurminen J, Luoma J, Taulu S, Ahonen A, Pekkonen E (2015b) Cortico-muscular coherence in advanced Parkinson’s disease with deep brain stimulation. Clin Neurophysiol 126:748–755.

Airaksinen K, Mäkelä JP, Taulu S, Ahonen A, Nurminen J, Schnitzler A, Pekkonen E (2011) Effects of DBS on auditory and somatosensory processing in Parkinson’s disease. Hum Brain Mapp 32:1091–1099.

Baker SN, Chiu M, Fetz EE, Fetz Afferent EE (2006) Afferent Encoding of Central Oscillations in the Monkey Arm. J Neurophysiol 95:3904–3910.

Baker SN, Olivier E, Lemon RN (1997) Coherent oscillations in monkey motor cortex and hand muscle EMG show task-dependent modulation. J Physiol 501:225–241.

Brown P, Oliviero a, Mazzone P, Insola a, Tonali P, Di Lazzaro V (2001) Dopamine dependency of oscillations between subthalamic nucleus and pallidum in Parkinson’s disease. J Neurosci 21:1033–1038.

Cheyne D, Ferrari P (2013) MEG studies of motor cortex gamma oscillations: evidence for a gamma “fingerprint” in the brain? Front Hum Neurosci 7:575.

Crone NE, Miglioretti DL, Gordon B, Lesser RP (1998) Functional mapping of human sensorimotor cortex with electrocorticographic spectral analysis. II. Event-related synchronization in the gamma band. Brain 121:2301–2315.

de Hemptinne C, Ryapolova-Webb ES, Air EL, Garcia P a, Miller KJ, Ojemann JG, Ostrem JL, Galifianakis NB, Starr P a (2013) Exaggerated phase-amplitude coupling in the primary motor cortex in Parkinson disease. Proc Natl Acad Sci U S A 110:4780–4785.

Deuschl G et al. (2006) A randomized trial of deep-brain stimulation for Parkinson’s disease. N Engl J Med 355:896–908.

Fernandez-Miranda JC, Pathak S, Engh J, Jarbo K, Verstynen T, Yeh FC, Wang Y, Mintz A, Boada F, Schneider W, Friedlander R (2012) High-definition fiber tractography of the human brain: Neuroanatomical validation and neurosurgical applications. Neurosurgery 71:430–453.

Gao W-J, Goldman-Rakic PS (2003) Selective modulation of excitatory and inhibitory microcircuits by dopamine. Proc Natl Acad Sci U S A 100:2836–2841.

Gao W-J, Wang Y, Goldman-Rakic PS (2003) Dopamine modulation of perisomatic and peridendritic inhibition in prefrontal cortex. J Neurosci 23:1622–1630.

Gradinaru V, Mogri M, Thompson KR, Henderson JM, Deisseroth K (2009) Optical deconstruction of parkinsonian neural circuitry. Science 324:354–359.

Gramfort A, Luessi M, Larson E, Engemann DA, Strohmeier D, Brodbeck C, Goj R, Jas M, Brooks T, Parkkonen L, Hämäläinen M (2013) MEG and EEG data analysis with MNE-Python. Front Neurosci 7:267.

Gross J, Kujala J, Hamalainen M, Timmermann L, Schnitzler A, Salmelin R (2001) Dynamic imaging of coherent sources: Studying neural interactions in the human brain. Proc Natl Acad Sci U S A 98:694–699.

Gwin JT, Ferris DP (2012) Beta- and gamma-range human lower limb corticomuscular coherence. Front Hum Neurosci 6:1–6.

Hammond C, Bergman H, Brown P (2007) Pathological synchronization in Parkinson’s disease: networks, models and treatments. Trends Neurosci 30:357–364.

Hirschmann J, Özkurt TE, Butz M, Homburger M, Elben S, Hartmann CJ, Vesper J, Wojtecki L, Schnitzler A (2013) Differential modulation of STN-cortical and cortico-muscular coherence by movement and levodopa in Parkinson’s disease. Neuroimage 68:203–213.

Hughes AJ, Daniel SE, Kilford L, Lees AJ, Daniel SE (1992) Accuracy of clinical diagnosis of idiopathic Parkinson’s disease: a clinico-pathological study of 100 cases. Neurosurgery, and Psychiatry 55:181–184.

Johnsen EL, Mogensen PH, Sunde NA, Østergaard K (2009) Improved asymmetry of gait in Parkinson’s disease with DBS: Gait and postural instability in Parkinson’s disease treated with bilateral deep brain stimulation in the subthalamic nucleus. Mov Disord 24:590–597.

Kang G, Lowery MM (2014) Effects of antidromic and orthodromic activation of STN afferent axons during DBS in Parkinson’s disease: a simulation study. Front Comput Neurosci 8:32.

Kass RE, Raftery AE (1995) Bayes Factors. J Am Stat Assoc 90:773–795.

Kilner JM, Baker SN, Salenius S, Hari R, Lemon RN (2000) Human cortical muscle coherence is directly related to specific motor parameters. J Neurosci 20:8838–8845.

Kristeva R, Patino L, Omlor W (2007) Beta-range cortical motor spectral power and corticomuscular coherence as a mechanism for effective corticospinal interaction during steady-state motor output. Neuroimage 36:785–792.

Kühn A a., Kupsch A, Schneider GH, Brown P (2006) Reduction in subthalamic 8-35 Hz oscillatory activity correlates with clinical improvement in Parkinson’s disease. Eur J Neurosci 23:1956–1960.

Lang AE, Lozano AM (1998) Parkinson’s Disease. N Engl J Med 339:1044–1053.

Limousin P, Krack P, Pollak P, Benazzouz A, Ardouin C, Hoffmann D, Benabid A-L (1998) Electrical Stimulation of the Subthalamic Nucleus in Advanced Parkinson’s Disease. N Engl J Med 339:1105–1111.

Litvak V, Eusebio A, Jha A, Oostenveld R, Barnes G, Foltynie T, Limousin P, Zrinzo L, Hariz MI, Friston K, Brown P (2012) Movement-Related Changes in Local and Long-Range Synchronization in Parkinson’s Disease Revealed by Simultaneous Magnetoencephalography and Intracranial Recordings. J Neurosci 32:10541–10553.

Mamikonyan E, Siderowf AD, Duda JE, Potenza MN, Horn S, Stern MB, Weintraub D (2008) Long-term follow-up of impulse control disorders in Parkinson’s disease. Mov Disord 23:75–80.

Manning JR, Jacobs J, Fried I, Kahana MJ (2009) Broadband shifts in local field potential power spectra are correlated with single-neuron spiking in humans. J Neurosci 29:13613–13620.

Marsden JF, Werhahn KJ, Ashby P, Rothwell J, Noachtar S, Brown P (2000) Organization of cortical activities related to movement in humans. J Neurosci 20:2307–2314.

Mattout J, Henson RN, Friston KJ (2007) Canonical source reconstruction for MEG. Comput Intell Neurosci 2007.

McIntyre CC, Hahn PJ (2010) Network perspectives on the mechanisms of deep brain stimulation. Neurobiol Dis 38:329–337.

McIntyre CC, Savasta M, Kerkerian-Le Goff L, Vitek JL (2004) Uncovering the mechanism(s) of action of deep brain stimulation: Activation, inhibition, or both. Clin Neurophysiol 115:1239–1248.

Medvedovsky M, Taulu S, Bikmullina R, Ahonen A, Paetau R (2009) Fine tuning the correlation limit of spatio-temporal signal space separation for magnetoencephalography. J Neurosci Methods 177:203–211.

Miller KJ, Zanos S, Fetz EE, den Nijs M, Ojemann JG (2009) Decoupling the cortical power spectrum reveals real-time representation of individual finger movements in humans. J Neurosci 29:3132–3137.

Mitra PP, Pesaran B (1999) Analysis of dynamic brain imaging data. Biophys J 76:691–708.

Morey RD, Rouder JN, Jamil T (2015) BayesFactor: Computation of Bayes Factors for Common Design. R package.

Muthukumaraswamy SD (2010) Functional properties of human primary motor cortex gamma oscillations. J Neurophysiol 104:2873–2885.

Nambu A, Tachibana Y (2014) Mechanism of parkinsonian neuronal oscillations in the primate basal ganglia: some considerations based on our recent work. Front Syst Neurosci 8:74.

Nambu A, Tokuno H, Hamada I, Kita H, Imanishi M, Akazawa T, Ikeuchi Y, Hasegawa N (2000) Excitatory cortical inputs to pallidal neurons via the subthalamic nucleus in the monkey. J Neurophysiol 84:289–300.

Nambu A, Tokuno H, Takada M (2002) Functional significance of the cortico- subthalamo- pallidal ‘hyperdirect’ pathway. Neuro Res 43:111–117.

Nolte G (2003) The magnetic lead field theorem in the quasi-static approximation and its use for magnetoencephalography forward calculation in realistic volume conductors. Phys Med Biol 48:3637–3652.

Oostenveld R, Fries P, Maris E, Schoffelen J-M (2011) FieldTrip: Open Source Software for Advanced Analysis of MEG, EEG, and Invasive Electrophysiological Data. Comput Intell Neurosci 2011:1–9.

Østergaard K, Aa Sunde N (2006) Evolution of Parkinson’s disease during 4 years of bilateral deep brain stimulation of the subthalamic nucleus. Mov Disord 21:624–631.

Park H, Kim JS, Paek SH, Jeon BS, Lee JY, Chung CK (2009) Cortico-muscular coherence increases with tremor improvement after deep brain stimulation in Parkinson’s disease. Neuroreport 20:1444–1449.

Petersen M V., Lund TE, Sunde N, Frandsen J, Rosendal F, Juul N, Østergaard K (2017) Probabilistic versus deterministic tractography for delineation of the cortico-subthalamic hyperdirect pathway in patients with Parkinson disease selected for deep brain stimulation. J Neurosurg 126:1657–1668.

Pollok B, Krause V, Martsch W, Wach C, Schnitzler A, Südmeyer M (2012) Motor-cortical oscillations in early stages of Parkinson’s disease. J Physiol 590:3203–3212.

R Core Team (2016) R: A Language and Environment for Statistical Computing.

Riddle CN, Baker SN (2005) Manipulation of peripheral neural feedback loops alters human corticomuscular coherence. J Physiol 566:625–639.

Salenius S, Avikainen S, Kaakkola S, Hari R, Brown P (2002) Defective cortical drive to muscle in Parkinson’s disease and its improvement with levodopa. Brain 125:491–500.

Salenius S, Portin K, Kajola M, Salmelin R, Hari R (1997) Cortical control of human motoneuron firing during isometric contraction. J Neurophysiol 77:3401–3405.

Schnitzler A, Gross J (2005) Normal and pathological oscillatory communication in the brain. Nat Rev Neurosci 6:285–296.

Shimamoto S a, Ryapolova-Webb ES, Ostrem JL, Galifianakis NB, Miller KJ, Starr P a (2013) Subthalamic nucleus neurons are synchronized to primary motor cortex local field potentials in Parkinson’s disease. J Neurosci 33:7220–7233.

Sridharan KS, Højlund A, Johnsen EL, Sunde NA, Johansen LG, Beniczky S, Østergaard K (2017) Differentiated effects of deep brain stimulation and medication on somatosensory processing in Parkinson’s disease. Clin Neurophysiol 128:1327–1336.

Surmeier DJ, Ding J, Day M, Wang Z, Shen W (2007) D1 and D2 dopamine-receptor modulation of striatal glutamatergic signaling in striatal medium spiny neurons. Trends Neurosci 30:228–235.

Taulu S, Hari R (2009) Removal of magnetoencephalographic artifacts with temporal signal-space separation: demonstration with single-trial auditory-evoked responses. Hum Brain Mapp 30:1524–1534.

Taulu S, Simola J (2006) Spatiotemporal signal space separation method for rejecting nearby interference in MEG measurements. Phys Med Biol 51:1759–1768.

Taulu S, Simola J, Kajola M (2005) Applications of the signal space separation method. IEEE Trans Signal Process 53:3359–3372.

Temperli P, Ghika J, Villemure J-G, Burkhard PR, Bogousslavsky J, Vingerhoets FJG (2003) How do parkinsonian signs return after discontinuation of subthalamic DBS? Neurology 60:78–81.

Tzourio-Mazoyer N, Landeau B, Papathanassiou D, Crivello F, Etard O, Delcroix N, Mazoyer B, Joliot M (2002) Automated anatomical labeling of activations in SPM using a macroscopic anatomical parcellation of the MNI MRI single-subject brain. Neuroimage 15:273–289.

van Doorn J, Ly A, Marsman M, Wagenmakers E-J (2016) Bayesian Inference for Kendall’s Rank Correlation Coefficient. Am Stat:0–0.

van Wijk BCM, Beek PJ, Daffertshofer A (2012) Neural synchrony within the motor system: what have we learned so far? Front Hum Neurosci 6:252.

Van Wijk BCM, Litvak V, Friston KJ, Daffertshofer a. (2013) Nonlinear coupling between occipital and motor cortex during motor imagery: A dynamic causal modeling study. Neuroimage 71:104–113.

Whitmer D, de Solages C, Hill B, Yu H, Henderson JM, Bronte-Stewart H (2012) High frequency deep brain stimulation attenuates subthalamic and cortical rhythms in Parkinson’s disease. Front Hum Neurosci 6:1–18.

Witham CL, Riddle CN, Baker MR, Baker SN (2011) Contributions of descending and ascending pathways to corticomuscular coherence in humans. J Physiol 589:3789–3800.

Witte M, Patino L, Andrykiewicz A, Hepp-Reymond MC, Kristeva R (2007) Modulation of human corticomuscular beta-range coherence with low-level static forces. Eur J Neurosci 26:3564–3570.

