## Supplementary figures and images for "Electromagnetic mapping of the effects of deep brain stimulation and dopaminergic medication on movement-related cortical activity and corticomuscular coherence in Parkinson’s disease"

### Supp. Fig. S1: DBS artefact rejection

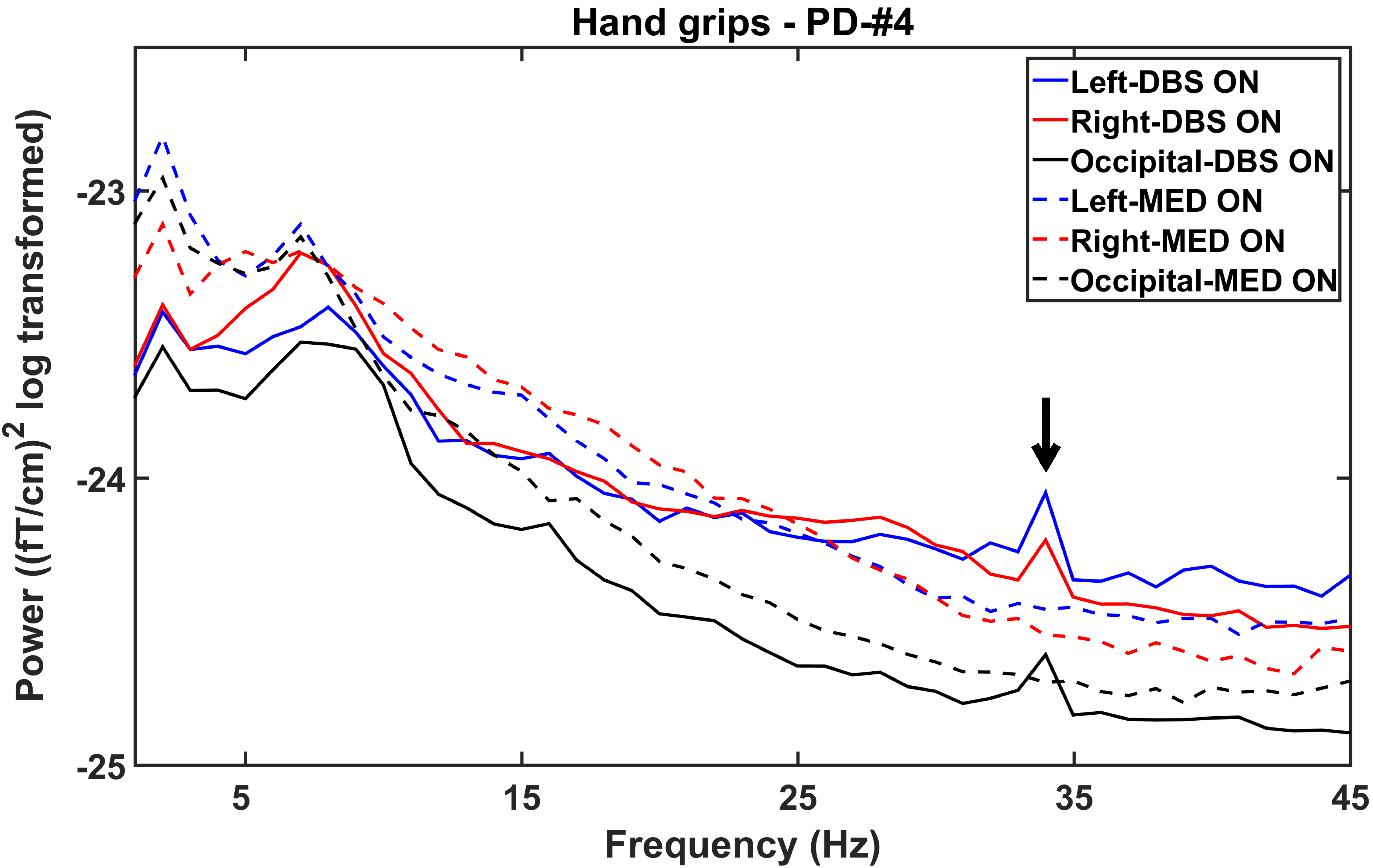

### Supp. Fig. S2: Correlation between beta-CMC and clinical motor state

#### Supp. Fig. S1 - Correlation- Beta-CMC vs Clinical motor state - MED ON condition

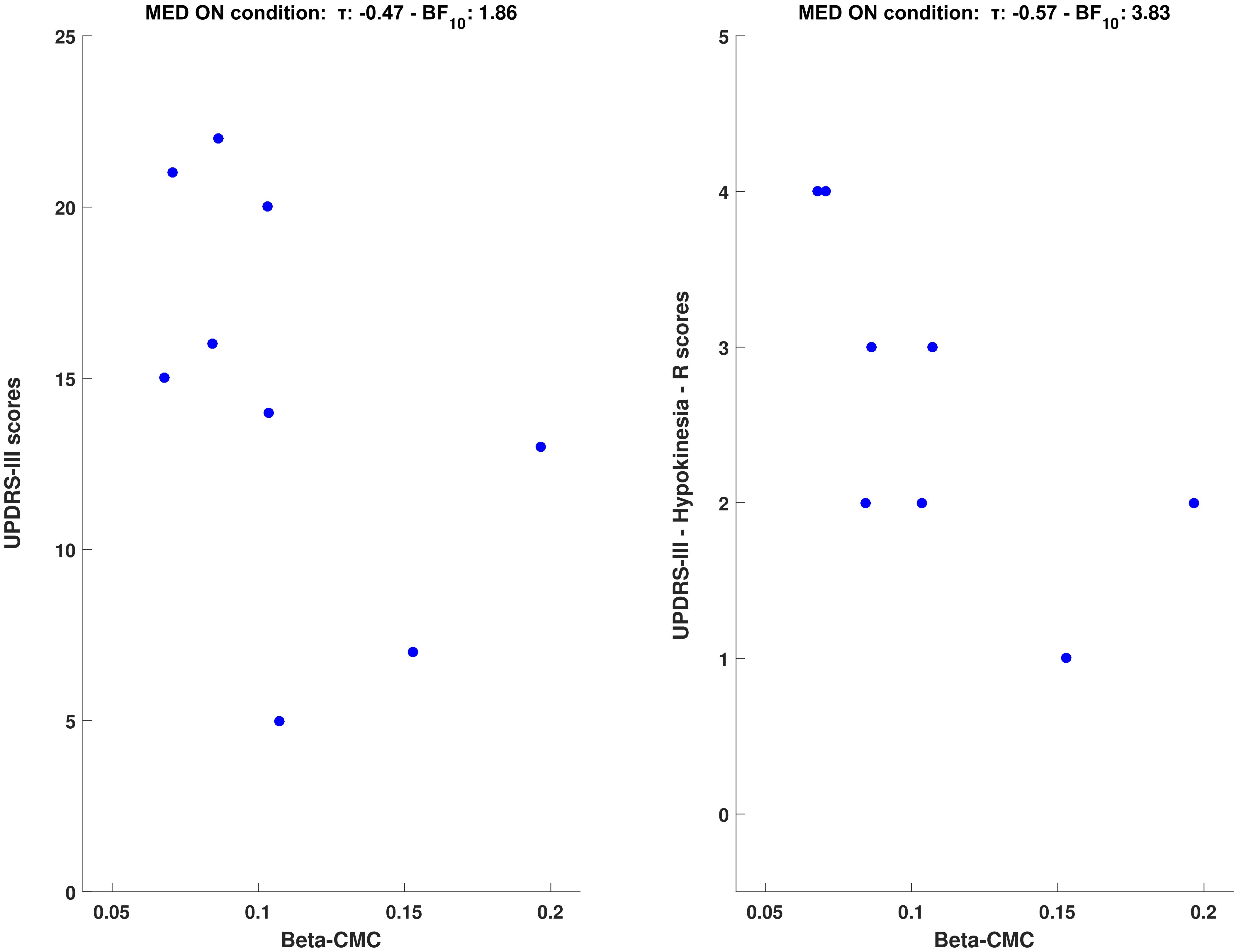

### Supp. Fig. S3: Influence of DBS artefacts on low-gamma power

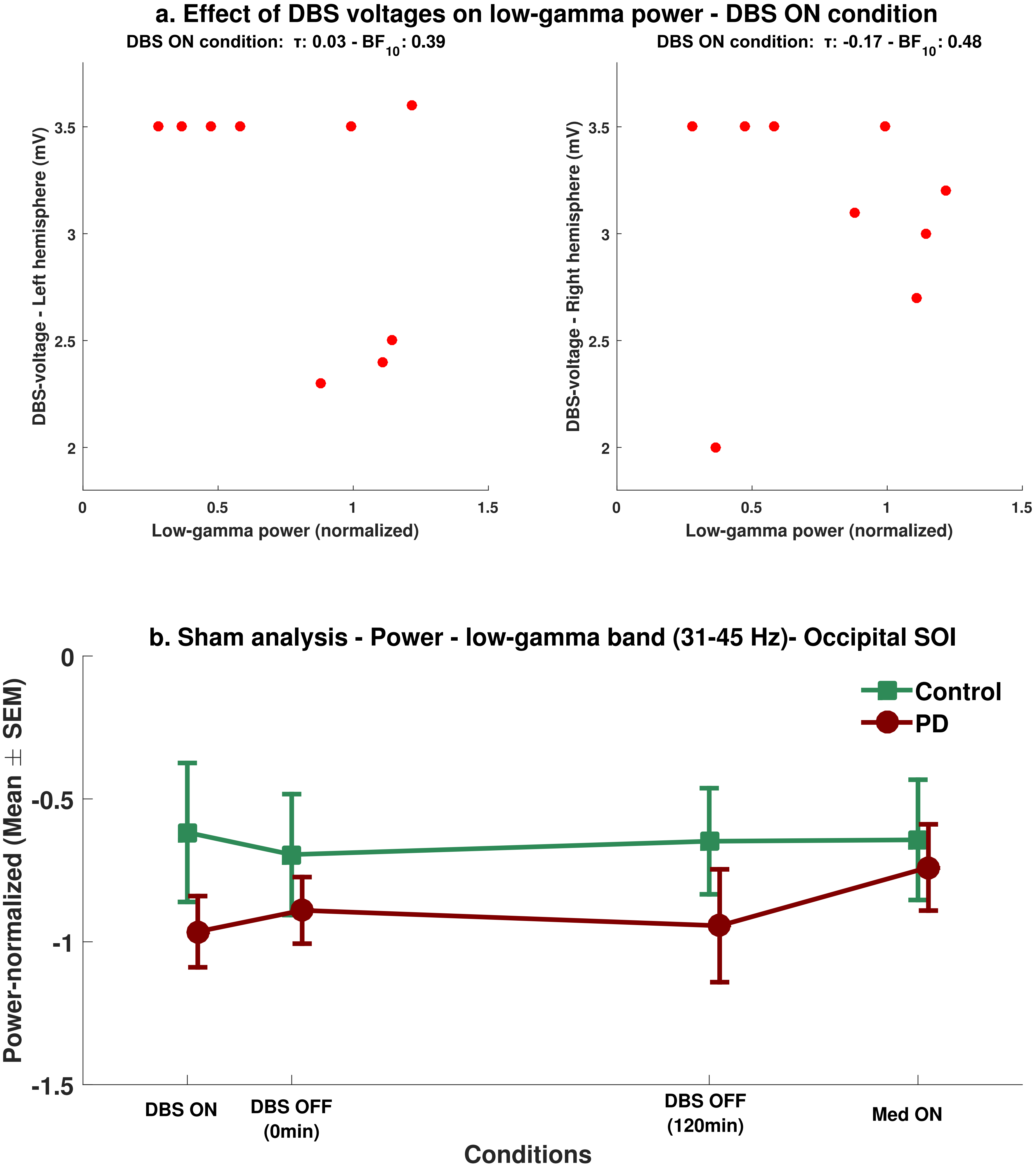
