## Supplementary material for "Electromagnetic mapping of the effects of deep brain stimulation and dopaminergic medication on movement-related cortical activity and corticomuscular coherence in Parkinson’s disease": Supp. Tab. ST1: Bayes Factors for the sham analyses

| $BF_{X0}$ -> alternative/null | | PD | | | | | | Ctrl | | | | | |
|---|---|---|---|---|---|---|---|---|---|---|---|---|---|
| | | All conditions ($H_1/H_0$) | 1 ≠ 2=3 ≠ 4 ($H_2/H_0$) | Specific hypotheses | | | | All conditions ($H_1/H_0$) | 1 ≠ 2=3 ≠ 4 ($H_2/H_0$) | Specific hypotheses | | | |
| | | | | 1:3 ≠ 4 ($H_3/H_0$) | 1 ≠ 2:4 ($H_4/H_0$) | 1=4 ≠ 2=3 ($H_5/H_0$) | 1=4 ≠ 2≠3 ($H_6/H_0$) | | | 1:3 ≠ 4 ($H_3/H_0$) | 1 ≠ 2:4 ($H_4/H_0$) | 1=4 ≠ 2=3 ($H_5/H_0$) | 1=4 ≠ 2≠3 ($H_6/H_0$) |
| **Right hemisphere** (Power) | **Beta** | **0.1926** | | | | | | **0.1698** | | | | | |
| | **Low-gamma** | **0.3306** | | | | | | **5.112** | | | | | |
| **Occipital** (Power) | **Beta** | **0.1365** | | | | | | **0.1396** | | | | | |
| | **Low-gamma** | **0.3352** | | | | | | **0.1725** | | | | | |
